# A neural hub that coordinates learned and innate courtship behaviors

**DOI:** 10.1101/2021.09.09.459618

**Authors:** Mor Ben-Tov, Fabiola Duarte, Richard Mooney

## Abstract

Holistic behaviors often require the coordination of innate and learned movements. The neural circuits that enable such coordination remain unknown. Here we identify a midbrain cell group (A11) that enables male zebra finches to coordinate their learned songs with various innate behaviors, including female-directed calling, orientation and pursuit. Anatomical mapping reveals that A11 is at the center of a complex network including the song premotor nucleus HVC as well as brainstem regions crucial to innate calling and locomotion. Notably, lesioning A11 terminals in HVC blocked female-directed singing, but did not interfere with female-directed calling, orientation or pursuit. In contrast, lesioning A11 cell bodies abolished all female-directed courtship behaviors. However, males with either type of lesion still produced songs when in social isolation. Lastly, monitoring A11 terminals in HVC showed that during courtship A11 inputs to the song premotor cortex signal the transition from innate to learned vocalizations. These results show how a brain region important to reproduction in both birds and mammals coordinates learned vocalizations with innate, ancestral courtship behaviors.

## Introduction

Complex appetitive behaviors, including foraging, hunting, tool use and courtship, often require the coordination of innate and learned movements (Demir et al., 2020; Keleman et al., 2012; Riotte-Lambert & Weimerskirch, 2013; Zampiga et al., 2006). For example, capuchin monkeys build on innate motor programs for grasping rocks and other objects to develop highly skilled and coordinated nut cracking skills (Ottoni & Izar, 2008), and male Drosophila (flies) learn to identify receptive mates by integrating innate and learned responses to female pheromones (Demir et al., 2020; Keleman et al., 2012). While studies in flies that have shown how learned and innate sensory cues are integrated in the brain to support successful courtship (Dickson, 2008; Keleman et al., 2012), little is known about how learned and innate motor programs are coordinated to enable complex courtship behaviors. The courtship display of the male zebra finch includes a learned song comprising a sequence of stereotyped syllables, or motif, that is seamlessly coordinated with a variety of innate behaviors, including female-directed calling, orientation, and pursuit (Klaus Immelmann, 1971; Ullrich et al., 2016; Williams, 2001) (Movie S1), providing the potential to understand the neural basis for such complex coordination. Despite this potential, little is known about the neural circuits that enable the coordinate of the learned and innate motor programs underlying the male zebra finch’s elaborate courtship behaviors.

Presumably, the neural circuits that enable such coordination must access motor regions for song as well as those that control innate vocalizations and female-directed head and body movements. Studies of zebra finches, as well as other songbirds, have thoroughly characterized how specialized forebrain nuclei, especially the song premotor nucleus HVC, control and pattern the motif (Appeltants et al., 2000; Aronov et al., 2008; Egger et al., 2020; Hahnloser et al., 2002; K. Hamaguchi & Mooney, 2012; Kosuke Hamaguchi et al., 2016; Long et al., 2010; Long & Fee, 2008; McCasland & Konishi, 1981; Nottebohm et al., 1976; Vu et al., 1998). In addition, vocal gating and locomotor circuits have been identified in the avian midbrain periaqueductal gray (PAG) (Fukushima & Aoki, 2000; Nieder & Mooney, 2020; Seller, 1981; Simpson & Vicario, 1990)) and pontine reticular formation (Steeves et al., 1987; D. M. S. Webster & Steeves, 1991; Deirdre M. S. Webster & Steeves, 1988), respectively. However, how activity is coordinated across such highly distributed motor regions to enable the male’s holistic courtship display remains a mystery. One idea is that a neural circuit that receives information about sexual motivation and communicates with HVC as well as the PAG and pontine reticular formation achieves this coordination.

The midbrain A11 cell group is part of a larger constellation of dopaminergic neurons found in all vertebrates (Anton Reiner et al., 1994; Smeets & González, 2000) that are implicated in motor control, motivation, reward and reproduction (Coddington & Dudman, 2018; da Silva et al., 2018; Goodson et al., 2009; Mohebi et al., 2019). The midbrain A11 in certain songbirds has been shown to receive input from the medial preoptic nucleus (POM) (Balthazart & Absil, 1997; Riters & Alger, 2004), a region important to appetitive and consummatory aspects of reproduction (Ball & Balthazart, 2004; Gahr, 2001; Trainor et al., 2003), and extends axons into HVC, raising the possibility that it functions to coordinate learned and innate aspects of the male’s courtship display. In fact, A11 neurons in male finches are activated in a variety of salient interactions with other zebra finches, including when an adult male engages in female-directed singing (Bharati & Goodson, 2006; Goodson et al., 2009), a sexually-motivated vocal display that is an essential appetitive component of successful courtship behavior in this songbird species. Here we combine pathway tracing combined with molecular phenotyping, cell- and axon-terminal specific manipulations, vocal analysis and machine learning tools, and in vivo calcium imaging, to reveal that A11 functions as a neural hub to smoothly and rapidly coordinate the learned and innate movements necessary to the complex courtship displays of male zebra finches.

## Results

### A11 axons target regions that encode learned and innate motor programs important to courtship

Although A11’s connections with HVC and POM have been previously described in songbirds, its broader pattern of afferent and efferent connectivity to other brain regions that may be important to courtship behaviors awaits description. To begin to provide such a description, we injected either conventional or genetically encoded anterograde tracers into the A11 of adult male zebra finches (Figure 1A, B Figure 1 – supplementary figure 1). These injections resulted in dense terminal labeling in the intercollicular nucleus of the dorsal midbrain (DM/ICo), which controls innate vocalizations in birds (Fukushima & Aoki, 2000; Seller, 1981; Simpson & Vicario, 1990) (Figure 1C); the gigantocellular part of the reticular nucleus of the caudal pons (RPgc), a major locomotor region (Steeves et al., 1987; D. M. S. Webster & Steeves, 1991; Deirdre M. S. Webster & Steeves, 1988) (Figure 1D); and also confirmed A11’s known projection to HVC (Appeltants et al., 2000; Ball & Balthazart, 2004) (Figure 1E). The locations of these three axonal terminations of A11 correspond to three elements of the male zebra finch’s holistic courtship display, namely female-directed innate vocalizations (DM/ICo), female pursuit and orientation (RPgc), and female-directed song (HVC). We also injected retrograde tracers into A11 of adult males to identify brain regions that provide it with synaptic inputs (Figure 1A). These injections identified three major sources of input to A11: the DM/ICo, the lateral deep cerebellar nucleus (lateral DCN, also known as the dentate nucleus), and the medial preoptic nucleus (POM) (Figure 1F-H). As such, A11 is situated to receive information about call generation (from the DM/ICo), sensorimotor integration and motor timing (through the DCN (Izawa et al., 2012; Proville et al., 2014)), and sexual motivation (via the POM (Ball & Balthazart, 2004; Balthazart & Absil, 1997; Gahr, 2001; Riters & Alger, 2004; Trainor et al., 2003)).

**Figure 1:**
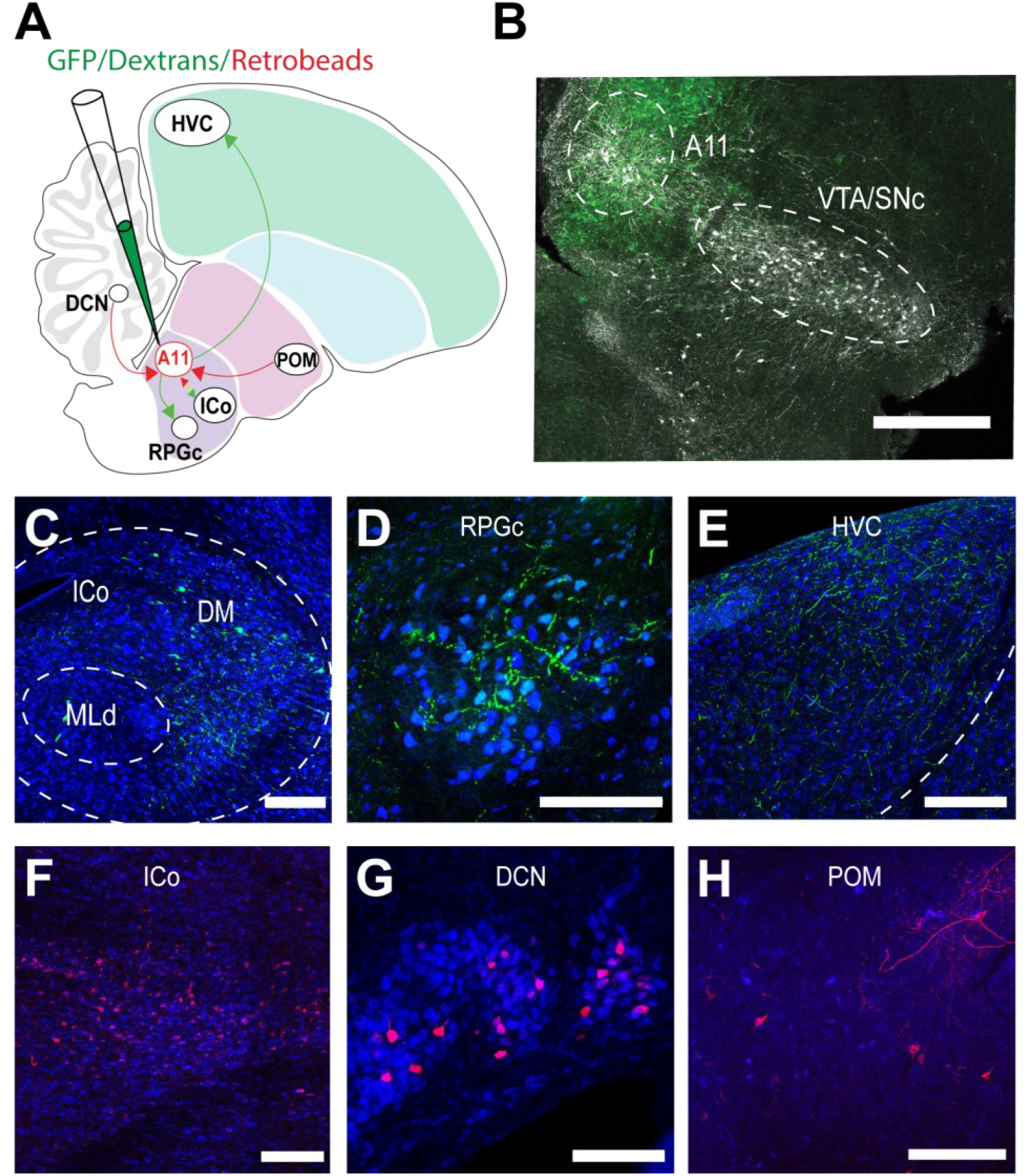
A11 axons target regions that encode learned and innate motor programs important to courtship. (**A**) Schematic of male finch brain in sagittal view showing injection of anterograde (GFP/dextrans) or retrograde (retrobeads/dextrans) tracers and A11’s efferents (green) and afferents (red). Consensus map is drawn from n=5 hemispheres from N=5 birds. (**B**) Representative injection site of a viral anterograde tracing strategy with GFP (green) and fluorescent antibody labeling of TH+ cells (pseudo-coloured grey). (**C-E**) Axonal projections from A11 to HVC, ICo and RPGc. (**G-H**) Cell bodies retrogradely labelled from A11 in DM ICo, DCN and POM. DCN, deep cerebellar nucleus; DM, dorsomedial part of the intercollicular nucleus (ICo); MLd, dorsal part of the mesencephalic nucleus; POM, medial preoptic nucleus; RPGc, gigantocellular part of the reticular nucleus of the caudal pons; SNc, substantia nigra pars compacta; VTA, ventral tegmental area. Scale bars in (C-H) are 200 μm. See also Figure 1 – figure supplement 1.

### A11 neurons synthesize both dopamine and glutamate

In addition to expressing TH, many DA neurons in mammals also express mRNAs for excitatory and inhibitory neurotransmitters, such as glutamate (Cai & Ford, 2018; Stuber et al., 2010) and gamma-aminobutyric acid (GABA) (Tritsch et al., 2012, 2014). To begin to better understand whether A11 cells in the zebra finch also might possess such dual transmitter identities, we combined retrograde tracing from HVC along with in situ hybridization chain reaction (Choi et al., 2018) for tyrosine hydroxylase (TH, a marker for dopaminergic neurons (Daubner et al., 2011)), the vesicular glutamate transporter 2 (VGLUT2, a marker of glutamatergic neurons (El Mestikawy et al., 2011)) and/or the vesicular inhibitory amino acid transporter (VGAT, a marker of GABA neurons (McIntire et al., 1997)). We found that the majority of TH+ A11 neurons, including those that project to HVC, were also VGLUT2+, while none were VGAT+ (Figure 2A-F). Thus, in addition to neuromodulatory effects mediated by DA, A11 neurons presumably also exert fast excitatory effects on their postsynaptic targets in HVC.

**Figure 2:**
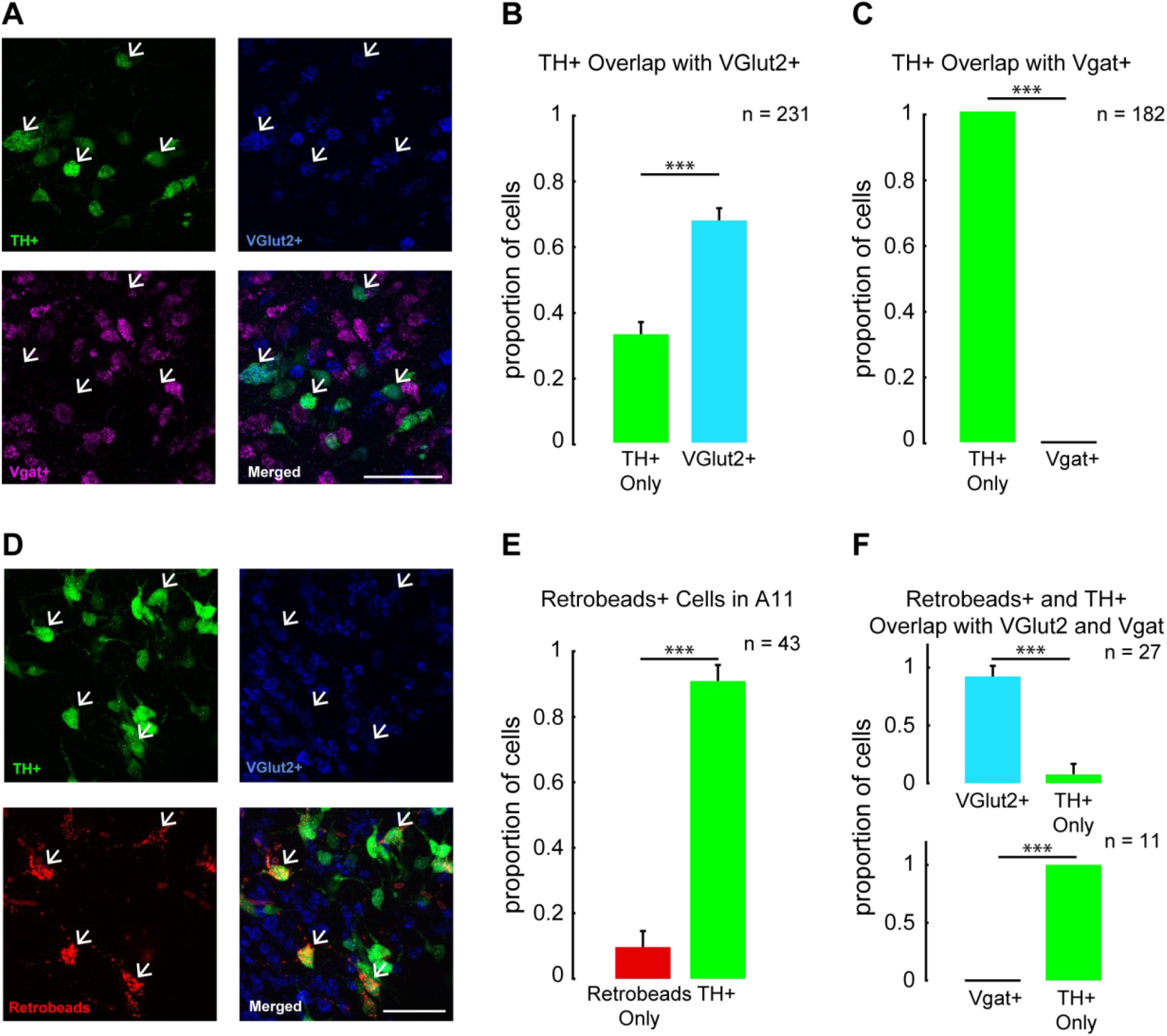
A11 neurons synthesize both dopamine and glutamate. (**A**) High-magnification images of A11 show the overlap of TH^+^ cells (green), VGlut2^+^ cells (blue) and VGAT^+^ cells (magenta). Scale bar, 100 μm. (**B**) Proportion of TH+ neurons in A11 that are also VGlut2^+^. χ^2^ test, χ^2^_1_=23.07, p<0.0001, n = 9 hemispheres from 5 birds. (**C**) Proportion of TH^+^ neurons in A11 that are also VGAT^+^. χ^2^ test, χ^2^_1_=182, p<0.0001, n = 5 hemispheres from 3 birds. (**D**) High-magnification images of A11 show the overlap of HVC-projecting neurons in HVC that are also TH^+^ cells (green) and VGlut2^+^ cells (blue). Scale bar, 100 μm. (**E**) Proportion of HVC-projecting cells in A11 that are also TH^+^. χ^2^ test, χ^2^_1_=25.33, p<0.0001, n = 6 hemispheres from 3 birds. (**F**) Proportion of TH^+^ HVC-projecting cells in A11 that are also VGlut2^+^ or VGAT^+^. χ^2^ test, χ^2^_1_=13.33, p=0.0002, n = 4 hemispheres from 2 birds and χ^2^_1_=11, p<0.0001, n = 2 hemispheres from 1 bird, respectively. Data are mean ± s.e.m.

### A11 cell bodies and their terminals in HVC are crucial for female-directed song, a learned behaviour

The divergent anatomical projections of A11 neurons raise the possibility that A11 plays an important role in male courtship behaviors. Therefore, we studied how the male behaved when placed in a chamber where visual access to the female could be carefully controlled via an electronic glass partition and monitored his vocal and non-vocal behaviors using audio and video recordings (Figure 3A). Notably, granting the male visual access to the female elicited a range of female-directed behaviors, including female directed singing (Figure 3B, (top panel)). Male zebra finches also sing in social isolation, a behavior known as undirected song, which is not sexually-motivated (Sossinka & Böhner, 1980). Males in our experimental chamber readily produced undirected song when visual access to the female was blocked (Figure 3B, (bottom panel)). We first compared each male’s singing behavior before and after we selectively lesioned either A11 terminals in HVC or A11 cell bodies in the midbrain with 6-hydroxydopamine (6-OHDA, (Hoffmann et al., 2016), Figure 3C,D,F,G and Figure 3 – supplementary figure 1A-E). Either treatment abolished female-directed singing in adult male zebra finches (Figure 3E,H). In several birds that were tracked for longer periods, this effect persisted for at least two months post-treatment. Surprisingly, male zebra finches with A11 terminal or cell body lesions recovered their ability to sing undirected songs, albeit at reduced rates, even though they failed to sing when presented with a female (Figure 3I-K). This recovery of undirected song was achieved within 5-10 days post-lesion. We also noted that syllable structure changed in males following A11 terminal lesions but remained intact following A11 cell body lesions (Figure 3 – supplementary figure 2A-I), which may reflect increased cell death in HVC that followed 6-OHDA treatment in this nucleus (Figure 3 – supplementary figure 2J). In summary, A11 plays a particularly important role in female-directed singing, the learned component of the male’s courtship display.

**Figure 3:**
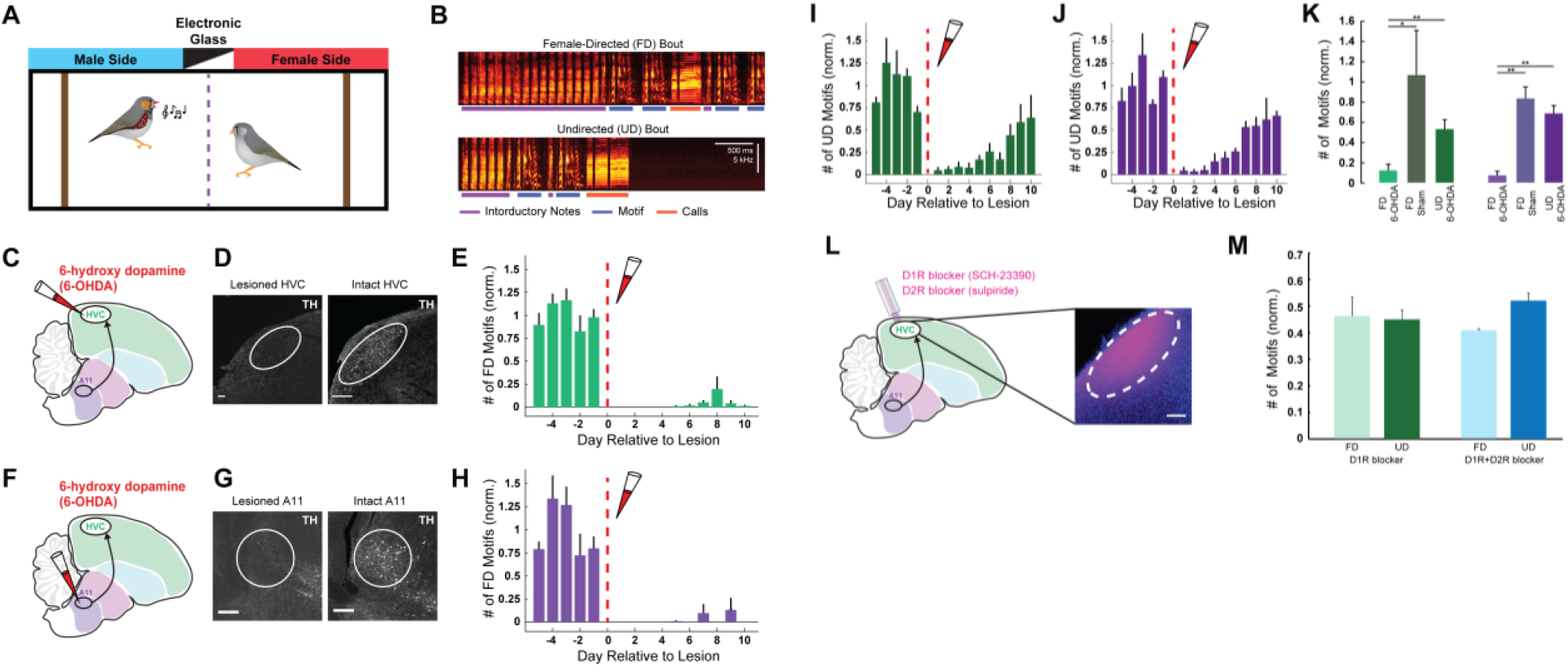
A11 cell bodies and terminals in HVC are crucial for learned female-directed vocalizations. (**A**) Schematic of the behavioral paradigm. (**B**) Example sonograms of female-directed (FD) song bout and undirected (UD) song bout. (**C**) Schematic of 6-OHDA injection to HVC, with the male finch brain shown in sagittal view. (**D**) Dopaminergic terminals in HVC labeled with TH antibody (pseudo-colored white) in a brain with 6-OHDA lesion in HVC (left) and an intact brain (right). (**E**) Mean normalized number of FD motifs before and after 6-OHDA injection in HVC (N=5 birds). (**F-H**) Same as (C-E) for N=4 birds treated with 6-OHDA in A11. (**I-J**) Same as (E and H) for undirected song bouts. (**K**) Mean post-treatment singing rates normalized to pre-treatment singing rates. Asterisks above horizontal bars indicate significant p values. UD singing rates were significantly higher than FD singing rates (paired t-test; p=0.0031 and p=0.0018 for HVC and A11 groups, respectively). FD singing rates were significantly higher for the control group than for the treated group (t-test; N=5 control birds and N=5 6-OHDA treated birds, p=0.034 and N=6 control birds and N=4 6-OHDA treated birds, p=0.0009 for HVC and A11 groups, respectively). Data are mean ± s.e.m. (**L**) Left: Schematic of the experiment, showing a microdialysis probe implanted in HVC and used to deliver D1 receptor (D1R) antagonist or D1R and D2 receptor (D2R) antagonist. Right: Confocal image showing that Fast Green (pseudo-colored magenta) diffused through the probe in HVC. Scale bar, 200 μm. (**M**) Number of female-directed and undirected motifs in D1R blockers days and D1R+D2R blockers days, normalized to saline days. In both contexts, DA receptor blockade decreases singing rates by approximately half. See also Figure 3 – figure supplements 1-2.

As just described, the A11 neurons that project to HVC synthesize and presumably release both DA and glutamate. To test whether DA release in HVC is the major driver of female-directed singing, we used microdialysis methods to reversibly block either D1-type or D1- and D2-type DA receptor signaling in the male’s HVC (Figure 3L). This treatment reduced both female-directed and undirected singing by about 50%, similar to levels of undirected singing observed in males treated with 6-OHDA in either HVC or A11 (Figure 3M). However, unlike these 6-OHDA males, which largely lost the ability to generate female-directed songs, males treated with D1 receptor blockers in HVC readily produced female-directed songs, albeit at a reduced rate (Figure 3M). This set of observations indicate that DA release from A11 axons in HVC facilitates singing regardless of social context, while also raising the possibility that glutamate released from A11 axons in HVC is necessary for female-directed singing.

### A11 cell bodies, but not A11 terminals in HVC, are important for innate courtship behaviors

We also examined these 6-OHDA-treated animals to determine whether A11 terminals in HVC or A11 cell bodies are important to innate aspects of the male’s courtship display (Figure 4). Males that were treated with 6-OHDA in HVC still produced a complete complement of innate courtship behaviors when granted visual access to a female. These innate behaviors included introductory notes, which are innate vocalizations that male zebra finches produce immediately prior to the learned song motif (Eales, 1985; Price, 1979; Sossinka & Böhner, 1980,). In fact, males treated with 6-OHDA in HVC produced prolonged strings of introductory notes but failed to produce a motif (Figure 4A (middle panel), B). These introductory notes were similar in their spectral structure to those produced prior to the 6-OHDA treatment (Figure 4 – supplementary figure 1A-D). Furthermore, performing movement analysis using a supervised learning algorithm (DeepLabCut) (Mathis et al., 2018) revealed that males pursued and oriented towards females in a similar manner before and after 6-OHDA treatment in HVC (Figure 4C left, D, F, G, Movie S2). In contrast, males treated with 6-OHDA in A11 lost all female-directed behaviors. Specifically, when presented with a female, males with A11 cell body lesions failed to produce introductory notes (Figure 4A (bottom panel), B), did not pursue or orient towards the female, and overall spent more time moving away from the female than they did prior to the lesion (Figure 4C-D, F, G, Movie S3). Notably, neither of these 6-OHDA treatments affected the male’s mobility, as the overall time males spent moving around the cage while the female was present was unchanged relative to pre-treatment levels (Figure 4E). Therefore, in adult male zebra finches, A11 plays an essential role in recruiting the learned song and various innate behaviors that characterize a holistic courtship display.

**Figure 4:**
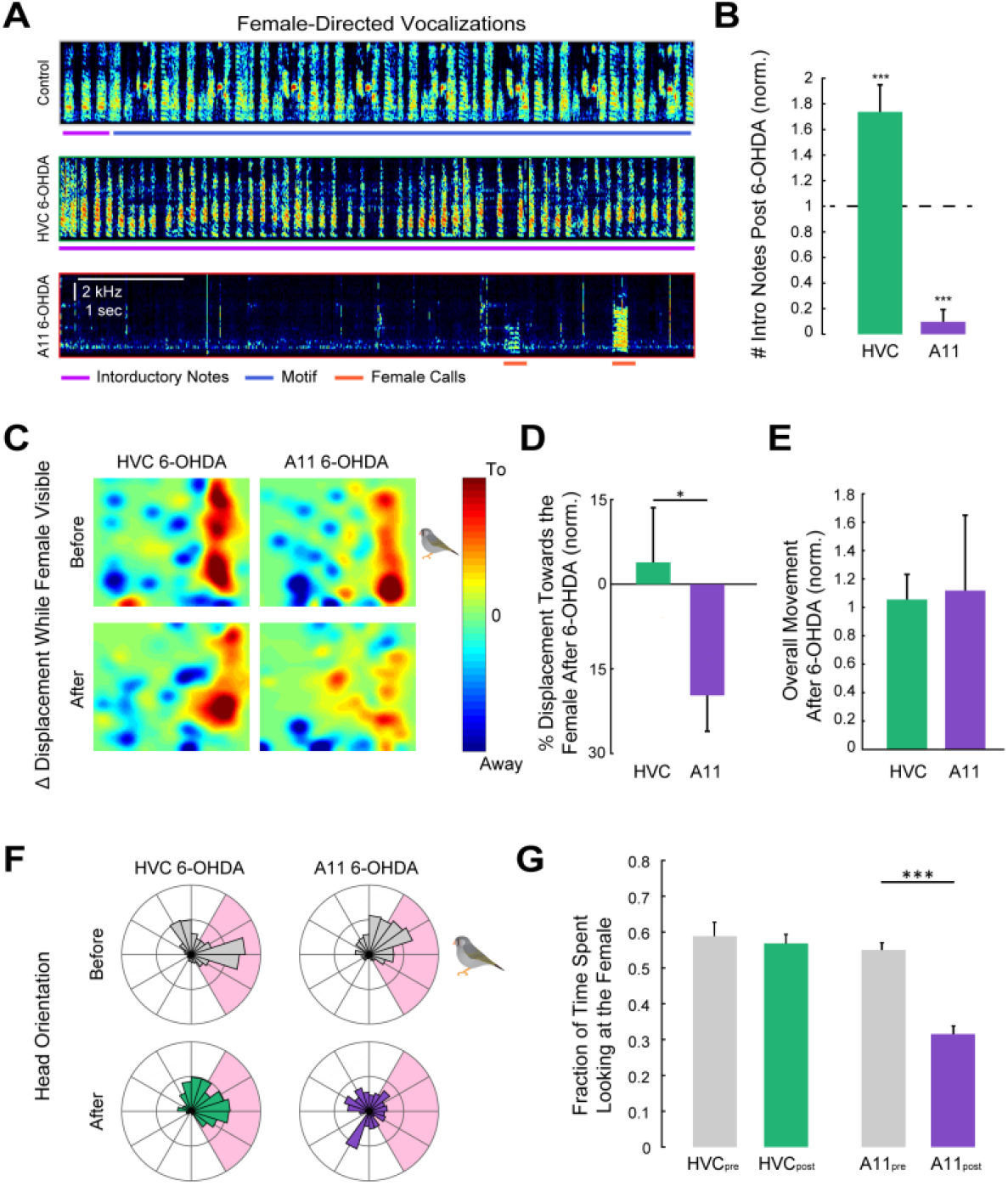
Activity of A11 cell bodies, but not their terminals in HVC, is important for innate courtship behavior. (A) Example sonograms of female-directed vocalizations for a control male, HVC 6-OHDA male and A11 6-OHDA male. Magenta, blue and red lines denote innate introductory, learned song motifs and female calls, respectively. (**B**) Mean normalized number of post-treatment FD introductory notes. The number of introductory notes produced significantly increased for the HVC 6-OHDA group and significantly decreased following A11 6-OHDA group (N=4 A11 treated birds and N=5 HVC treated birds, two-way repeated measures ANOVA p<0.0001, followed by Bonferroni’s multiple comparisons test, p=0.0018 and p=0.0012 for HVC and A11 groups, respectively). (**C**) Change in male’s displacement after female presentation, before and after 6-OHDA treatment. Warm and cool colors represent displacement towards and away from the female, respectively. (**D**) Mean displacement towards or away from the female post-treatment, normalized to pre-treatment values. HVC 6-OHDA males moved towards the female significantly more than A11 6-OHDA males (t-test, p=0.049). (**E**) Mean overall movement post-treatment, normalized to pre-treatment values. There was not a significant difference between the two groups (t-test; p=0.91). (**F**) Example of head orientation distributions after female presentation, before and after 6-OHDA treatment. Pink shading represents the angle range in which the female was visible to the male. (**G**) Fraction of the time males spent looking at the female pre- and post-treatment for the two experimental groups. A11 6-OHDA males significantly reduced the time they spent looking at the female (two-way repeated measures ANOVA p=0.0006. Post-hoc comparisons with Bonferoroni’s correction, for the A11 group, p=0.0001). Data are mean ± s.e.m. See also Figure 4 – figure supplement 1.

### Monitoring the activity of A11 terminals in HVC during directed and undirected song

Here we observed that males in which A11 terminals in HVC were lesioned could not produce female-directed songs but still generated introductory notes (Figure 4A). Furthermore, although HVC is not necessary for the production of innate vocalizations (Aronov et al., 2008; Nottebohm et al., 1976), which are instead gated and patterned by brainstem nuclei (Fukushima & Aoki, 2000; Seller, 1981; Simpson & Vicario, 1990; J. M. Wild et al., 1997; J. Martin Wild, 1993), prior studies have shown that HVC neurons exhibit elevated activity that is time-locked to production of introductory notes and other innate calls (Benichov et al., 2016; Benichov & Vallentin, 2020; Daliparthi et al., 2019; McCasland & Konishi, 1981; Yu & Margoliash, 1996). In light of these prior studies, our current observations raise the possibility that, during courtship, A11 transmits information about innate vocalizations to HVC that helps to coordinate the transition from introductory notes to the learned motif. To test this idea, we injected AAV axon-targeted GCaMP (Broussard et al., 2018) into the A11 of male zebra finches, waited several (3-8) weeks for expression of GCaMP in A11 axons in HVC (Figure 5 – supplementary figure 1) and then used fiber photometry to measure calcium signals in these A11 axons as the male interacted with a female partner (Figure 5A; Figure 5 – supplementary figure 2). These experiments revealed that, during female-directed singing, calcium signals in A11 axons in HVC were elevated prior to motif onset, during the period when the male is producing introductory notes (Figure 5B). Alignment of the calcium signal to the male’s vocal display showed that A11 axon activity started to increase above baseline well (> 1 sec) before the first introductory note and peaked at motif onset (Figure 5E-G, Figure 5 – supplementary figure 3A-B).

**Figure 5:**
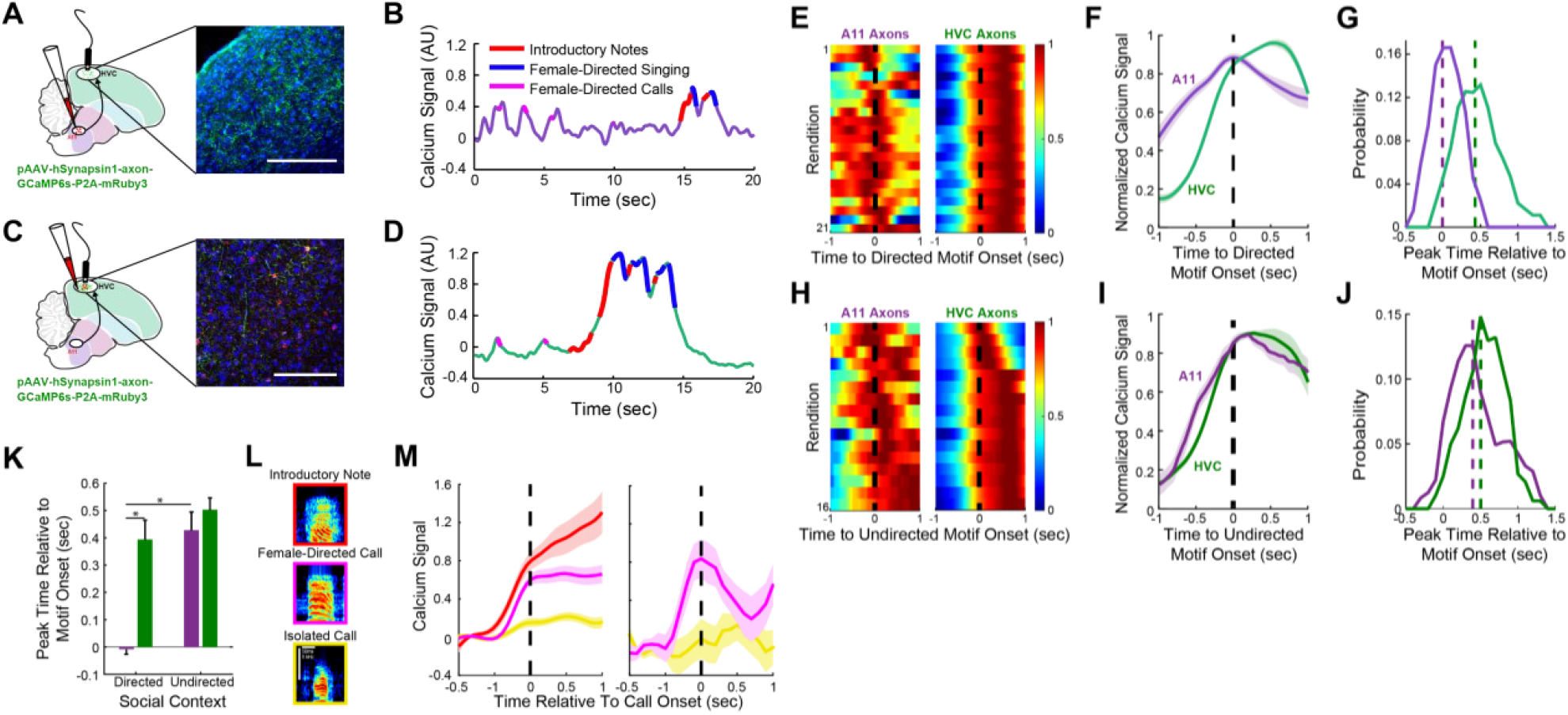
Vocalization-related activity of A11 axons in HVC depends on social context. (A) Left: schematic of male finch brain in sagittal view showing fiber photometry imaging of A11 axons in HVC (A11 axons) using axon-targeted GCaMP6s. Right: GFP+ A11 axons in HVC labeled in green. (B) Calcium signal of A11 axons in HVC of a male finch as he calls and sings to a female. Overlaid on the calcium signal are red, blue and magenta lines that denote the production of the male’s introductory notes, song motif and female-directed calls, respectively. (C, D) Same as (A, B) for axon-targeted GCaMP6s injection in HVC (HVC axons). mRuby+ cell bodies labeled in red and GFP+ axons labeled in green. (E) Example calcium traces during female-directed singing, aligned to motif onset (black dashed line), for one A11 axons bird and one HVC axons bird. (F) Mean normalized calcium activity during female-directed song motifs for the birds presented in (E). Black dashed line denotes the motif onset. (G) Peak activity time relative to motif onset. Purple and green dashed lines represent the mean peak activity of A11 axons and HVC axons, respectively (N=4 A11 axons birds and N=4 HVC axons birds). (H-J) Same as (E-G), for undirected song motifs. Data in (H-I) are from the same birds as in (E-F). Data in (J) are from N=3 A11 terminals birds and N=3 HVC axons birds. (K) Mean peak activity latencies for A11 axons (purple bars) and HVC axons (green bars) in female-directed and undirected singing relative to motif onsets. Peak activity of A11 axons during female-directed singing is significantly earlier than HVC axons peak activity during female-directed singing and from A11 axons activity during undirected singing (mixed-effects analysis, p<0.05. p<0.0005 and p<0.01 for the two comparisons, respectively). (L) Example sonograms of the three types of the male’s innate vocalizations: introductory notes (red), female-directed calls (magenta) and isolated calls (yellow) (M) Baseline subtracted calcium signals of A11 axons during innate vocalizations in two males. Data are mean ± s.e.m of 12-90 isolated calls. Activity during female-directed vocalizations was significantly higher than during isolated vocalizations (t-test with Holm–Bonferroni correction for multiple comparisons, p<0.001). See also Figure 5 – figure supplements 1-4.

A similar imaging strategy in which we injected AAV axon-targeted GCaMP directly into HVC (Figure 5C, D) was used to detect elevations in vocalization-related calcium signals in local HVC axons. These experiments revealed that vocalization-related increases in calcium signals in local HVC axons were delayed relative to A11 axon activity, with local axonal activity increasing above baseline ∼0.5 seconds before the first introductory note and peaking in mid-motif (Figure 5E-G, Figure 5 – supplementary figure 3A-B). In contrast, during undirected song, peaks in A11 and HVC axon activity were almost simultaneous, a temporal difference from directed song that was accounted for by a rightward shift in A11 axon activity relative to motif onset (Figure 5H-J). Therefore, while A11 axons in HVC are active during both directed and undirected singing, as expected given that HVC and A11 are both nodes within a recurrent network (K. Hamaguchi & Mooney, 2012), A11 leads HVC activity during directed song whereas these two regions are simultaneously active during undirected song (Figure 5K). In addition to these timing differences, higher levels of activity in A11 also characterize female-directed singing, as expression of the immediate early gene c-fos was higher during female-directed relative to undirected song (Figure 5 – supplementary figure 4A-C), in agreement with an earlier study (Bharati & Goodson, 2006). Lastly, the calcium-related activity of A11 axons in HVC also increased when the male produced introductory notes that did not lead to a motif and during innate calls that were female-directed, but not when the male called in social isolation (Figure 5L, M). Taken together with our lesion studies, these imaging results support a sequential process in which A11 signals HVC about the generation of female-directed innate vocalizations, such as calls and introductory notes, helping to coordinate the transition to the learned song motif.

## Discussion

To our knowledge, this study provides the first evidence of a neural hub that coordinates learned and innate motor programs to generate a holistic suite of courtship behaviors. As a variety of skilled appetitive behaviors in birds and mammals, including foraging, hunting, and tool use, also depend on the coordination of learned and innate movements (Demir et al., 2020; Keleman et al., 2012; Ottoni & Izar, 2008; Riotte-Lambert & Weimerskirch, 2013; Zampiga et al., 2006), the present findings may point to a general strategy by which midbrain cell groups participate to organize these highly skilled behaviors. Effectively, our results provide a motor counterpart to studies in flies that have shown how learned and innate sensory cues are integrated in the brain to support successful courtship (Demir et al., 2020; Dickson, 2008).

A striking feature of the sexually-motivated courtship display in male zebra finches is the seamless and rapid transition from innate motor programs, including those for female-directed orientation, pursuit and calling, to the learned motor program that patterns the motif. The anatomical and functional studies undertaken here illuminate how the midbrain A11 can act as a central hub to enable such seamless and rapid motor integration (schematized in Figure 6). To successfully court a female, the male produces song bouts comprising extended and tightly interwoven sequences of innate introductory notes and learned song syllables, generated with millisecond precision over many seconds (Woolley & Doupe, 2008). The present fiber photometry findings point to the ascending projections from A11 to HVC as a major source of information about female-directed innate vocalizations, presumably mediated by its input from call production centers in the ICo (Figure 1F), that help to precisely time the transition between innate and learned vocalizations and enhance the potency of sexual signaling. While A11 is active during both female-directed and undirected singing, A11 terminal activity in HVC peaks earlier relative to motif onset and c-fos expression in A11 cell bodies is more elevated during female-directed song. This shift in the timing and magnitude of A11 activity parallels a shift in A11’s functional influence on song, as determined by 6-OHDA lesions and DA receptor microdialysis experiments, with A11 playing an obligatory, leading role to drive directed singing and merely a facilitatory role in undirected singing. More generally, because A11 and HVC are part of a recurrent network (Hamaguchi & Mooney, 2012), our findings may point to how song production shifts from bottom-up control during encounters with a female to top-down control when the male vocalizes in social isolation.

**Figure 6:**
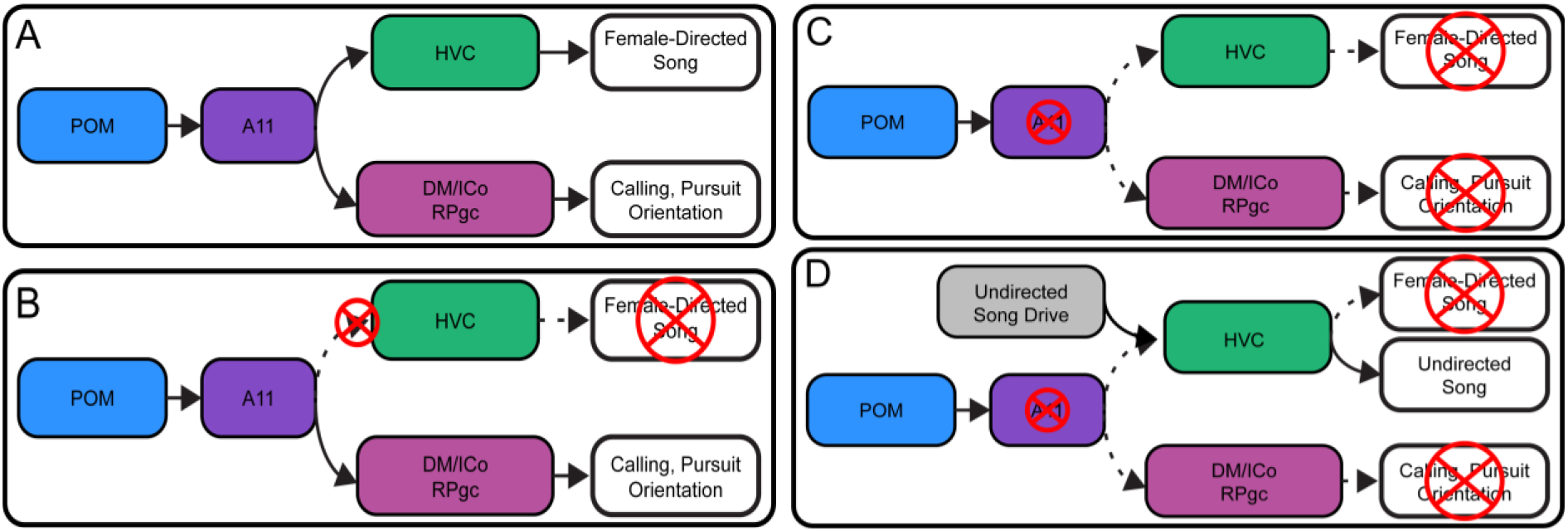
A schematic describing the circuit for the coordination of learned and innate courtship behavior in male zebra finch. (**A**) A11 receives projections from arousal and reproductive centers (POM) and in return sends projections to the song premotor nucleus HVC and center controlling innate calling and locomotion, DM/ICo and RPgc, respectively. These projections enable the male to sing, call, orient and pursue the female. (**B**) When we abolish A11 projections to HVC, the male is unable to produce female-directed learned songs, while the innate courtship behaviors are intact. (**C**) When we abolish A11 cell bodies, the male is unable to produce both learned and innate elements of courtship behavior. (**D**) However, males can still produce isolated, undirected songs, which implies the existence of undirected song drive to HVC.

Prior studies have detected that the often-subtle acoustic features that distinguish female-directed song from undirected song greatly enhance its salience as a sexual signal (Houtman, 1992; Woolley & Doupe, 2008). A reasonable idea is that these acoustic differences can be attributed to the male’s sexual motivation and arousal (Riters et al., 2000). As suggested by a recent study in canaries (Haakenson et al., 2020), the projections from the POM to A11 provide a plausible pathway for sexual motivation and arousal signals to reach forebrain song circuitry, while also accounting for the elevated A11 activity that occurs in adult males producing female-directed songs (Figure 5 – supplementary figure 4A-C and Bharati & Goodson, 2006). Moreover, our finding of the selective role that A11’s projections to HVC play in triggering the directed but not undirected song motif (Figure 3) underscores that the circuitry that promotes these two types of singing is at least partially distinct (Walters et al., 1991).

The midbrain A11 cell group in mammals, the likely homologue of the songbird midbrain A11 (Anton Reiner et al., 1994; Kingsbury et al., 2011), is implicated in various innate appetitive and consummatory aspects of reproduction through its descending projections to motor centers in the brainstem and spinal cord (Giuliano & Allard, 2001). The present results show how this ancestral reproductive structure has expanded to incorporate a learned behavior into the male’s holistic courtship display via its ascending projections to song premotor circuitry in the forebrain. The evolutionarily ancient nature of A11 raises the possibility that it serves a similar function in other species that depend on such complex motor integration during courtship, including our own.

## Materials and Methods

### Data reporting

No statistical methods were used to predetermine sample size. The experiments were not randomized and the investigators were not blinded to allocation during experiments and outcome assessment.

### Animal model

Adult males and adult females (>105 days old) zebra finches (Taeniopygia guttata) were obtained from the Duke University Medical Center breeding facility. All experimental procedures were in accordance with the NIH guidelines and approved by the Duke University Medical Center Animal Care and Use Committee. Birds were kept under a 14:10-hr light:dark cycle with free access to food and water. Data were collected from 50 adult male zebra finches.

### Tissue collection

Birds were deeply anesthetized with intramuscular injection of 20 μl Euthasol (Virbac), and transcardially perfused with 0.025 M phosphate-buffered saline (PBS) followed by 4% paraformaldehyde (PFA). Brains were removed, post-fixed in 4% PFA at 4°C overnight and moved to cryoprotective 30% sucrose PFA solution for two days. Frozen sagittal sections (thickness of 50 or 75 μm) were prepared with a sledge microtome (Reichert) and collected in PBS.

### Immunofluorescence

Floating sections were washed three times in PBS, permeabilized with 0.3% Triton X-100 in PBS (PBST) for 10 minutes and incubated with rabbit primary antibody against TH (1:1000, AB152; Millipore/Sigma) at 4°C overnight. Sections were then washed three times in PBS and incubated with anti-rabbit secondary antibody (1:500; Jackson ImmunoResearch) in PBS at room temperature for 2-4 h, followed by three washes in PB. Sections were coverslipped with Fluoromount-G (SouthernBiotech), and then imaged with a confocal microscope (Zeiss) through a 20x or 10x objective lens controlled by Zen software (Zeiss). To label A11 projections and inputs, AAV2/9.CAG-scGFP (made in lab), dextran (Alexa Fluor 488, D-22910, ThermoFisher) or retrobeads (LumaFluor) were injected into the A11 of adult male birds 4–7 days before perfusion. Images were processed with ImageJ to adjust for contrast. For the analysis of TH^+^ neurons in A11 after 6-OHDA treatments, neurons with diameter greater than 10 µm were counted manually.

### Floating section *in situ* hybridization chain reaction (HCR)

Birds were perfused with 4%PFA/PBS and postfixed in the same solution overnight and then in 30% sucrose in DEPC-PBS for 2 overnights at 4°C. Brains were then sectioned at 75 µm and collected into 0.5-1% PFA (4% PFA diluted in RNAse-free PBS). At room temperature, slices were first washed twice in PBS for 3 min, incubated in 5% SDS/PBS for 45 min, rinsed twice with 2x sodium chloride sodium citrate 0.1% Tween 20 (2x SSCT), and then were put in 2x SSCT for 15 min on a shaker. Then they were hybridized in 2.5 µL probe set/500 µl probe hybridization buffer overnight at 37°C. The probes were custom made by Molecular Instruments to detect zebra finch isoforms of vGluT2, TH, and VGAT. The next day, slices were washed four times for 15 min with 500 µL of probe wash buffer at 37°C and twice in 2x SSCT for 5 min at room temperature on a shaker. Then they were incubated in 500 µl of HCR amplification buffer for 30 min at room temperature on a shaker. Last, slices were incubated in a solution containing 300 µL HCR amplification buffer and fluorescent hairpins for the HCR initiator probe for 2 overnights, in the dark at 25°C. On the last day, at room temperature, slices were washed twice with 2x SSCT for 5 min, stained with Neurotrace for 2 hours (1:500, N21479; Invitrogen), rinsed twice with 2x SSCT and mounted on a slide with Fluoromount-G. Hairpins, probe sets and probe hybridization buffer were created by Molecular Instruments. HCR for TH and vGluT2 was performed on sections from 4 birds, HCR for TH and VGAT was performed on sections from 2 birds, and HCR for TH, vGluT2 and VGAT was performed on sections from 1 bird.

### Lesion experiments

A pair of male and female adult zebra finches were housed in an isolated soundproof box. The male and the female were separated by electronic glass (HOHOFILM Electronic PDLC, Smart Film) that was connected to an external switch. When powered off, this glass is opaque, preventing the male and the female from seeing each other. In order to record female-directed singing, the experimenter powered the glass on, rendering it transparent and enabling the birds to see each other. The use of electronic glass eliminated the need to handle the birds, increasing the probability that the males would sing to the female. Video recordings started approximately 20 seconds prior to visibility onset. The glass remained transparent for 1-7 minutes. Video recordings stopped approximately 20 seconds after visibility offset. Baseline singing rates were recorded for 5-7 days, after which birds were divided into 4 experimental groups, HVC 6-OHDA, HVC sham, A11 6-OHDA, and A11 sham (Figure 1 K). Videos of the birds were recorded using webcams (Genius WideCam F100). Songs were automatically recorded with Sound Analysis Pro (SAP2011^33^). Singing rates were calculated manually by counting all female-directed songs produced during the first minute of female presentation and by counting all undirected songs produced during a 4-hour period each day. Female-directed and undirected song rates were calculated for five days prior to surgery. An average singing rate over this baseline period was calculated and then used to normalize singing rates for each day pre- and post-surgery. Vocalizations of >5 ms were detected by amplitude thresholding of the recorded sounds. Pairwise similarity scores (asymmetrical similarity) between either song motifs or introductory notes from pre-treatment days and post-treatment days were calculated using SAP2011^33^ with default parameters for zebra finches.

### Terminal deoxynucleotidyl transferase-mediated dUTP nick-end labeling (TUNEL) assay

Birds lesioned in HVC with 6-OHDA were perfused, and brain sections were collected as described in the previous sections. Detection of apoptosis was carried out with some modifications to the DeadEnd™ Colorimetric TUNEL System (Promega). Briefly, free-floating sections were fixed in 4% PFA for 15 mins., and washed three times with PBS at room temperature. The sections were digested in a Proteinase K solution (20ug/ml) for 10-15 mins., followed by three PBS washes and an additional fixation step. The samples were placed on a shaker at room temperature in an equilibration buffer for 10 mins., followed by an incubation step with a biotinylated nucleotide mix and rTdT enzyme in a 37°C humidified chamber for 1 hour. The sections were washed once with 2xSSC for 15 mins., and three times with PBS. Then, they were incubated for 60 mins. at room temperature with AlexaFluor 488- streptavidin (1:500, ThermoFisher) and washed 3 times with PB before mounting.

### Video analysis

Videos were analyzed for 20 seconds prior to female presentation and 30 seconds after female presentation. For each video, the bird’s position and head orientation were measured using either DeepLabCut (Mathis et al., 2018) or custom MATLAB graphical user interfaces (M. Ben-Tov) that enable the marking of the bird’s body position and head orientation across video frames. For each video, the position was normalized to the cage size, to allow a comparison between different videos and birds.

### General surgery

Adult male birds were anesthetized with 1%–2% isoflurane inhalation and placed in a stereotaxic device on a heat blanket. Stereotaxic coordinates and multiunit recordings were used to localize target sites for injection and implantation. Stereotaxic coordinates, measured from the bifurcation of the midsagittal sinus, were 0.0 mm rostral, 2.4 mm lateral and 0.5 mm ventral (head angle of 18°) for HVC and 3.4 mm rostral, 0.7 mm lateral and 6.1 mm ventral (head angle of 62°) or 0.35 caudal, 0.7 mm lateral and 5.15 ventral (head angle of 22°) for A11. Reagents (Dextran, Alexa Fluor 488, 594 or 647, Invitorgen; RetroBeads 590, Lumafluor) or viruses were injected using Nanoject-II (Drummond Scientific). Viral injections were performed bilaterally with a volume of 300–1000 nL per hemisphere. Viruses and plasmids were obtained from the Penn Vector Core (Pennsylvania, USA) and the UNC Vector Core (Chapel Hill, USA).

### Injection of 6-OHDA

Adult male birds received bilateral injections of either 400nl 6-OHDA solution into HVC or 80-100nl 6-OHDA solution into A11 (N=4 for A11 and N=5 HVC). The solution was PBS-based and included 10-60 mM 6-OHDA hydrochloride (Tocris, 2547), 10 μM l-ascorbic acid (Millipore/Sigma, A92902), and 1 μM desipramine hydrochloride (Tocris, 3067), which was included as an inhibitor for noradrenaline and serotonin transporters to protect noradrenergic and serotonergic neuron terminals at the injection site. Control birds received an injection of PBS with 10 μM ascorbic acid and 1 μM desipramine (N=6 for A11 sham group and N=5 for HVC sham group). Drugs were dissolved in PBS immediately before injection in place of equimolar NaCl (working solution: ∼300 mOsm, pH 7.3). After injection, birds were returned to their original home cage until approximately 14 days post injection.

### Microdialysis infusion of drugs

Adult male birds were implanted bilaterally with custom-made microdialysis probes and then housed individually in an acoustic sound-proof box. Beginning two days later, saline or dopamine blocker were infused on alternate days into HVC at light onset (D1-type blocker: 5 mM R(+)-SCH-23390 hydrochloride, Millipore/Sigma, D054, D2-type blocker: 5 mM S-(-)- sulpiride, Tocris, 0895). Songs were recorded starting an hour after the infusion for three hours total. Female-directed songs were collected by introducing a female for one-minute sessions, 4-5 sessions per day. Song counts were calculated for 7 birds (D1-type blocker) and 2 birds (D1-type and D2-type blockers), in 3 saline days and 2 dopamine blockers days.

### Fiber photometry imaging

Adult male birds were injected with pAAV-hSynapsin1-axon-GCaMP6s-P2A-mRuby3 (axon-target GCaMP6s) bilaterally into A11 or HVC. After waiting a minimum of 3 weeks for viral expression, birds were anesthetized and placed in a stereotaxic apparatus. Bilateral craniotomies were made over HVC and fiber optic ferrules (200 um core, 0.37 NA, Neurophotometrics) were implanted. For all recordings, axon-targeted GCaMP6s was excited at two wavelengths (470nm for imaging of calcium-dependent signals and 415nm for an interleaved isosbestic control to eliminate motion artifacts). An sCMOS camera was used to capture fluorescence (FP3001, Neurophotometrics) at 30 Hz. Synchronized video and sound recordings were acquired using a webcam (Logitech). Data acquisition was performed with custom Bonsai code and data analyzed using custom-written Matlab scripts. In each imaging session, the signal from the isosbestic control channel was first smoothed and then regressed to the signal from the calcium-dependent channel. To calculate the calcium-dependent signal, first the linear model generated from regression was used to generate a predicted control signal. Then the calcium-dependent signal was calculated by subtracting the predicted control signal from the raw calcium-dependent signal (Figure. S3). Audio recordings were filtered using a third-order median filter. Introductory notes, syllables and different types of calls were labeled manually in Matlab. Calcium signals were aligned to the audio recordings and then z-scored to normalized changes in fluorescence across animals.

### Hybridization chain reaction for identification of Fos expression

In order to minimize off-target Fos mRNA detection, birds were perfused 30 minutes after cage lights first turned on in the morning. For female-directed singing (n = 4 birds), 4-5 females were presented sequentially to maximize motif amounts. Live video was monitored to ensure no significant undirected (facing away from the female, disengaged) singing occurred during female presentations. For the undirected condition (n = 4 birds), birds were allowed to sing freely in the morning for the 30-minute window. Lights were then turned off and birds were immediately perfused. Brains were processed for HCR as described in a previous section, the custom probes designed by Molecular Instruments targeted Fos, TH and VGAT. For each bird, a z-stack encompassing A11 was collected at 40x power to accurately visualize Fos signal, along with a TH channel. All image processing was done with ImageJ. First, the TH and Fos channels were noise-subtracted (20 μm rolling ball radius), and TH channel was smoothed, automatically thresholded (otsu method), and converted to a binarized mask. The mask was then transferred to the Fos channel, which was then also thresholded. Fos particles within the TH mask were then quantified for intensity (expressed as a fraction of the TH mask).

### Statistics

Data are shown as mean ± s.e.m., unless otherwise noted. One-way ANOVAs, two-ways ANOVAs, and their corresponding post-hoc comparisons were performed on Prism (GraphPad). T-tests were performed in Matlab. To examine the different proportion of labelled neurons in the A11, χ2 tests were performed (Figure 2B,C,E,F). A paired t-test was used to examine the effect of 6-OHDA in HVC and in A11 on the production of female-directed and undirected singing rates (Figure 3K). A t-test was used to compared female-directed singing between 6-OHDA and sham injections in HVC and in A11 (Figure 3K). Two-way repeated measurements ANOVA (p(Interaction)<0.0001, followed by Bonferroni’s multiple comparisons test was performed to examine 6-OHDA treatment effect on the production of introductory notes (Figure 4B). A t-test was used to test for a significant effect of 6-OHDA treatment in HVC or in A11 on the movement towards the female (Figure 4D) and on overall movement (Figure 4E).

One-way ANOVAs were performed to examine whether 6-OHDA injections abolished A11 neurons (F_2,9_ = 13.70, p=0.0019 and F_2,9_ = 9.18, p=0.0067; Figure 3 – supplement figure 1D and 1E). One-way repeated measurement ANOVAs were performed to examine the effect of 6-OHDA treatment on the pairwise similarities between song motifs and introductory notes for the HVC group and song motifs for the A11 group (for the HVC 6-OHDA song motifs: F_2,8_ = 6.751, p<0.05, followed by post-hoc Tukey test, asterisk denotes p<0.05; Figure 3 – supplement figure 2I and Figure 4 – supplement figure 1D). A mixed-effect analysis was used to test for a significant difference in peak response time in calcium signals between A11 terminals in HVC and HVC local axons, in female-directed and undirected singing (mixed-effects analysis, p(interaction)=0.0226. p=0.0002 and p=0.011 for A11 axons and HVC axons in directed singing, and A11 axons in directed and undirected singing, respectively; Figure 5K). A t-test was used to test the rise time relative to first introductory note difference between the A11 axons group and the HVC axons group (Figure 5 – supplementary figure 3B). A t-test was used to test the ratio of Fos positive TH positive cells in A11 during female-directed and undirected song, p=0.0033 (Figure 5 – supplementary figure 4C).

## Acknowledgements

We thank Fan Wang, Kevin Franks, Masashi Tanaka, Katherine Tschida, Audrey Mercer and Thomas Pomberger for critical discussion and for reading earlier versions of this manuscript.

## Supplemental Information

**Figure 1 – supplementary figure 1.**
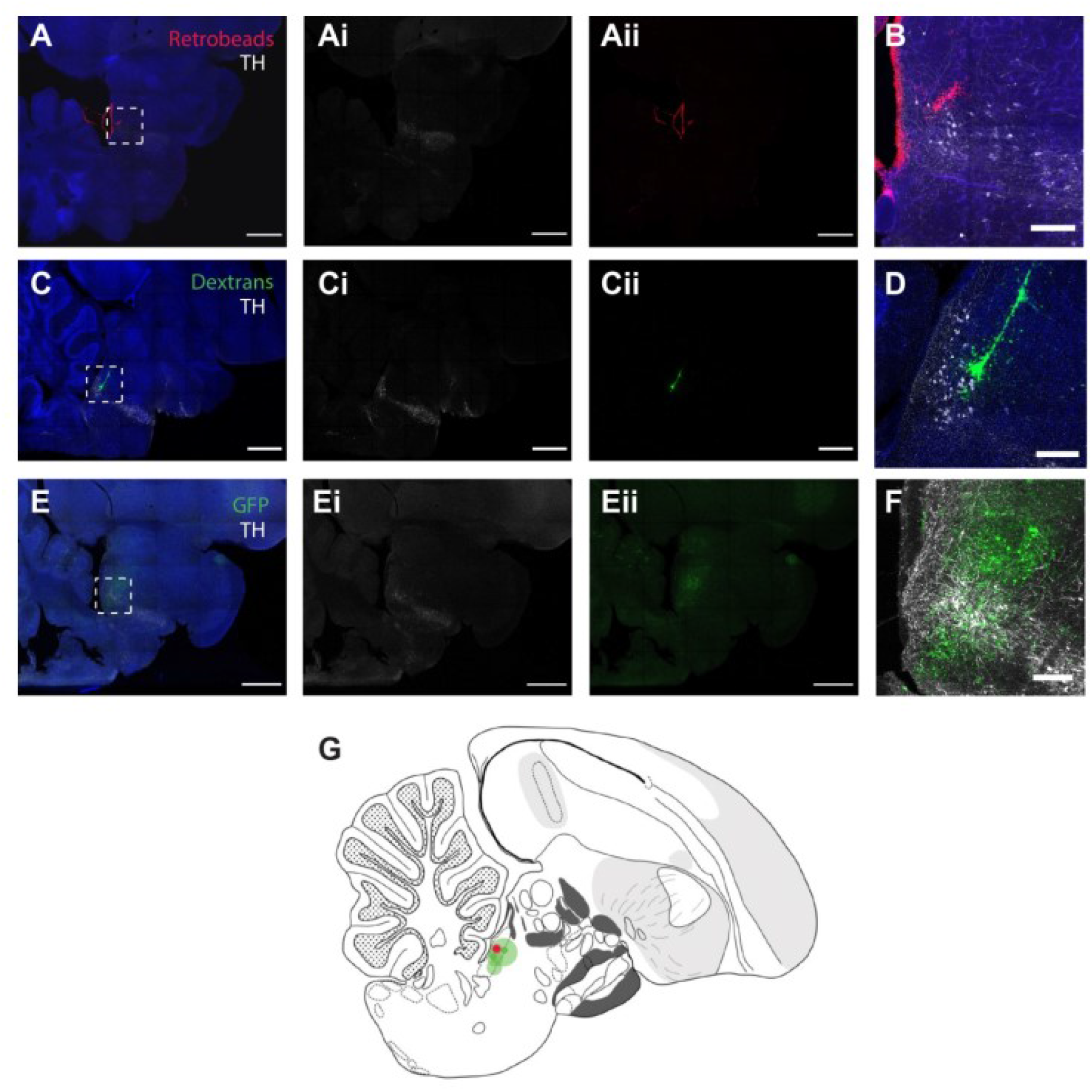
Tracing A11 inputs and outputs. (**A**) Representative sagittal section of the injection site in A11 for retrograde tracing (red) with fluorescent Nissl-staining (blue) and TH antibody labeling (pseudo-colored grey), scale bars, 1 mm. Left panel shows image composite and subsequent panels (ai,aii) individual channels. (**B**) area magnification of the rectangle shown in (a), scale bar, 200 μm. (**C-F**) Same as (A) and (B) for bidirectional tracing with dextrans (green) and anterograde viral tracing with GFP (green). (**G**) Schematic of injection sites and approximate diffusion of the tracers mapped on a sagittal section of the histological atlas (N = 5 hemispheres from 5 birds).

**Figure 3 – supplementary figure 1.**
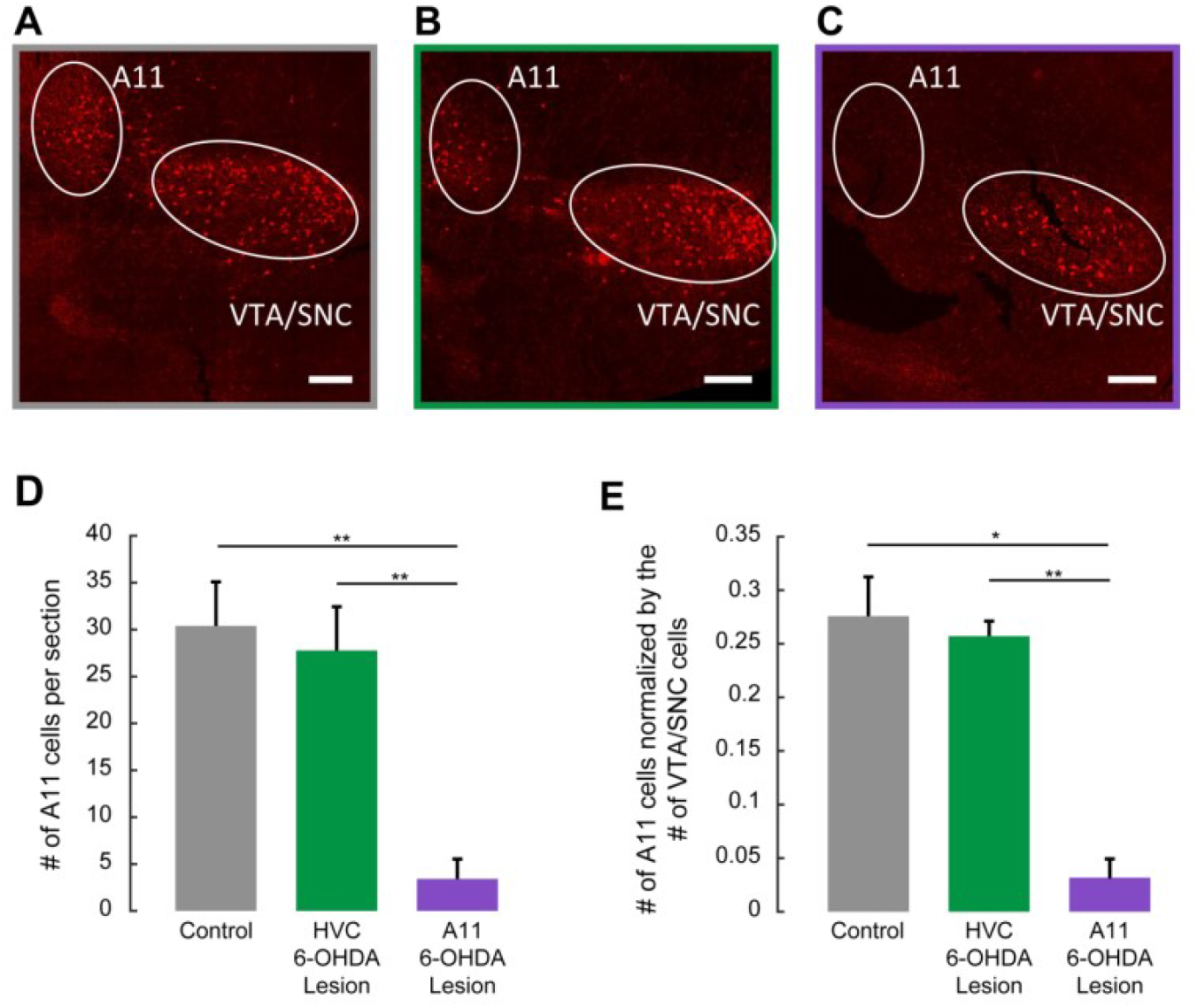
Effects of 6-OHDA injection into A11 or HVC on DA cell bodies in the midbrain. (**A**) A maximum-projected confocal image of serial sagittal sections, showing A11 and VTA/SNc TH immunolabelling (approximately 0.5 mm lateral of the midline) in control birds. Similar results were obtained in 4 independently repeated experiments. Scale bar is 200 μm. (**B**) A maximum-projected confocal image of serial sagittal sections, showing A11 and VTA/SNc TH immunolabelling in a bird that was injected with 6-OHDA in HVC. Similar results were obtained in 4 independently repeated experiments. Scale bar is 200 μm. (**C**) Same as (A), showing A11 and VTA/SNc TH immunolabelling, in a bird that was injected with 6-OHDA in A11. Similar results were obtained in 4 independently repeated experiments. Scale bar is 200 μm. (**D**) Mean number of A11 cells in maximum-projected sections for the three experimental groups (1-4 sections scored from N=4 birds in each condition). Number of A11 TH+ cells was significantly greater in control and HVC 6-OHDA groups than in the A11 6-OHDA group (one-way ANOVA, p<0.005, followed by post-hoc Tukey tests, two asterisks denote p<0.01). e, Number of A11 cells in a maximum-projected section normalized to the number of VTA/SNc cells from the same section for the three experimental groups. Number of normalized A11 TH+ cells were significantly lower in the A11-treated group than the number of A11 TH+ cells in the other groups (one-way anova, p<0.01, followed by post-hoc Tukey tests, one asterisk denotes p<0.05, two asterisks denote p<0.01).

**Figure 3 – supplementary figure 2.**
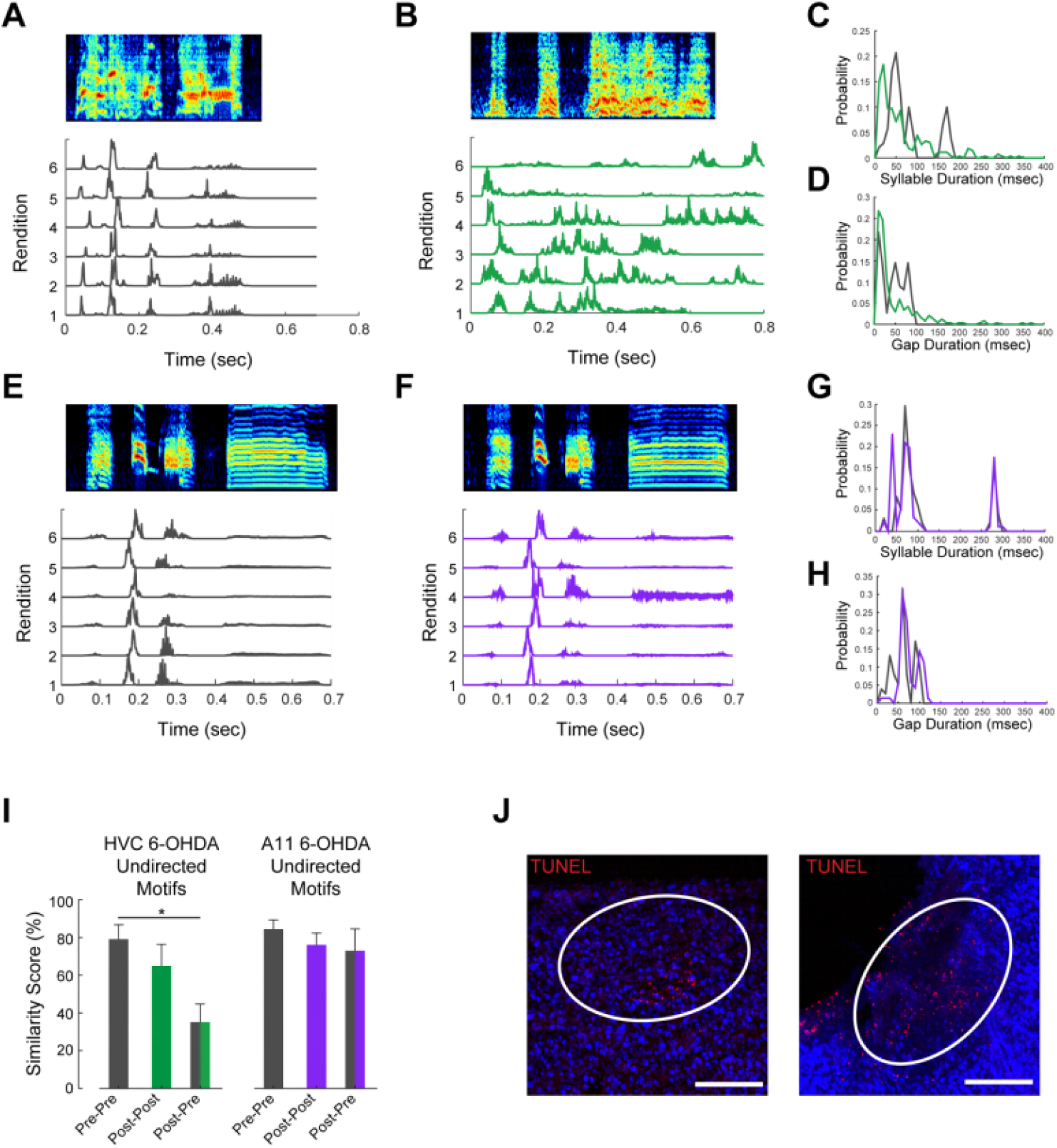
6-OHDA treatment effect on song motifs. (**A**) An example sonogram (top) and amplitude envelopes (bottom traces) of undirected song renditions of an intact, untreated male zebra finch. (**B**) Same as in (A), for the same male after treatment with 6-OHDA in HVC. (**C**) Distributions of syllable durations for pre-treatment (grey) and post HVC 6-OHDA treatment (green). (**D**) Distributions of inter-syllable gap durations for pre-treatment (grey) and post HVC 6-OHDA treatment (green). (**E-H**) Same as (A-D), for A11 6-OHDA treatment. (**I**). Pairwise similarity scores for randomly selected undirected motifs (left-most columns for HVC 6-OHDA treated birds, right-most columns for A11 6-OHDA treated birds). Data are mean ± s.e.m. Only the motifs produced by the HVC 6-OHDA treated group were significantly less similar post treatment compared to pre-treatment. (One-way repeated measurements ANOVA p<0.05. Post-hoc Tukey test, asterisk denotes p<0.05). (**J**) Two examples of terminal deoxynucleotidyl transferase dUTP nick end labeling (TUNEL), used to detect DNA breaks formed during the final phase of apoptosis, after the injection of 6-OHDA in HVC, scale bars 200 μm.

**Figure 4 – supplementary figure 1.**
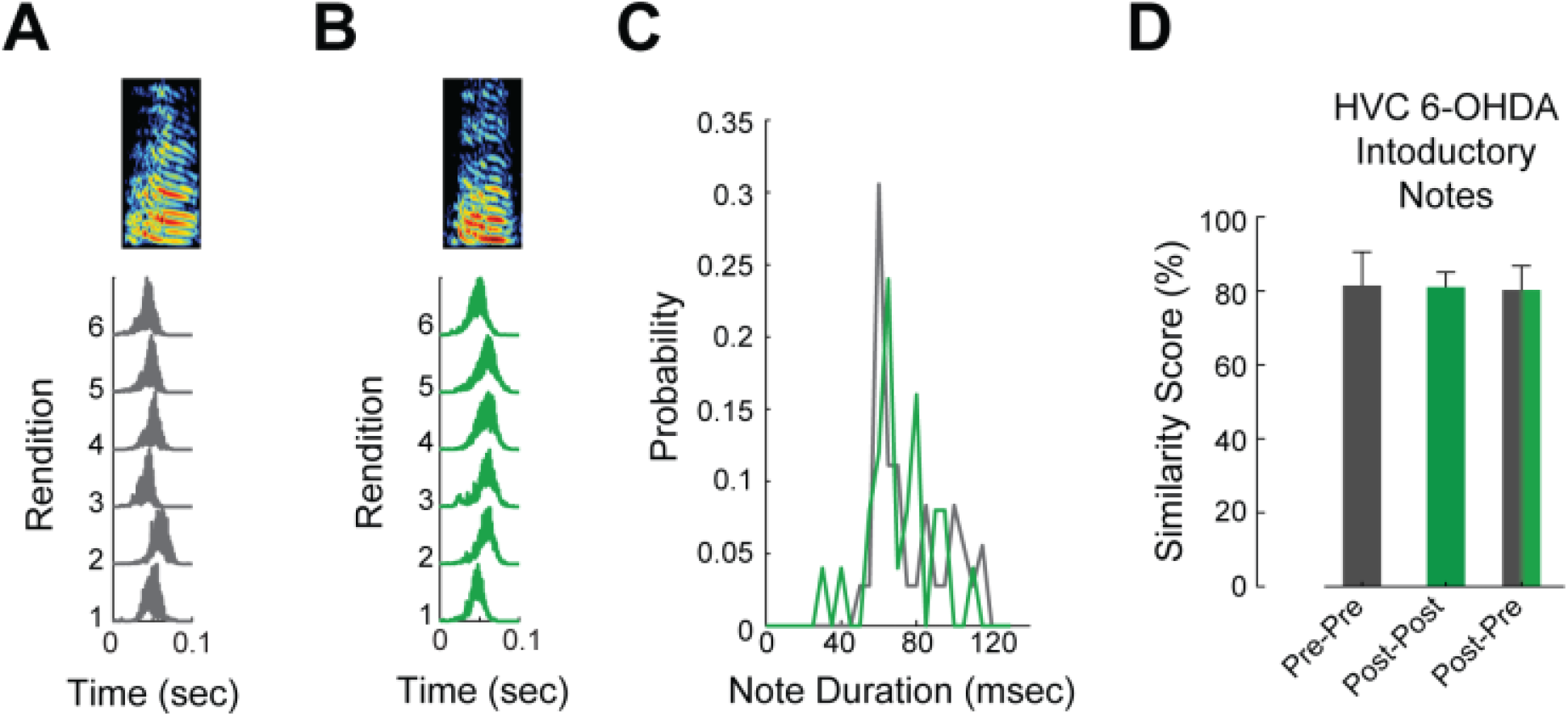
6-OHDA treatment effect on introductory notes. (**A**) An example sonogram (top) and amplitude envelopes (bottom traces) of introductory notes of an intact, untreated male zebra finch. (**B**) Same as in (A), for the same male after treatment with 6-OHDA in HVC. (**C**) Distributions of introductory note durations for pre-treatment (grey) and post HVC 6-OHDA treatment (green). (**D**) Pairwise similarity scores for randomly selected introductory notes.

**Figure 5 – supplementary figure 1.**
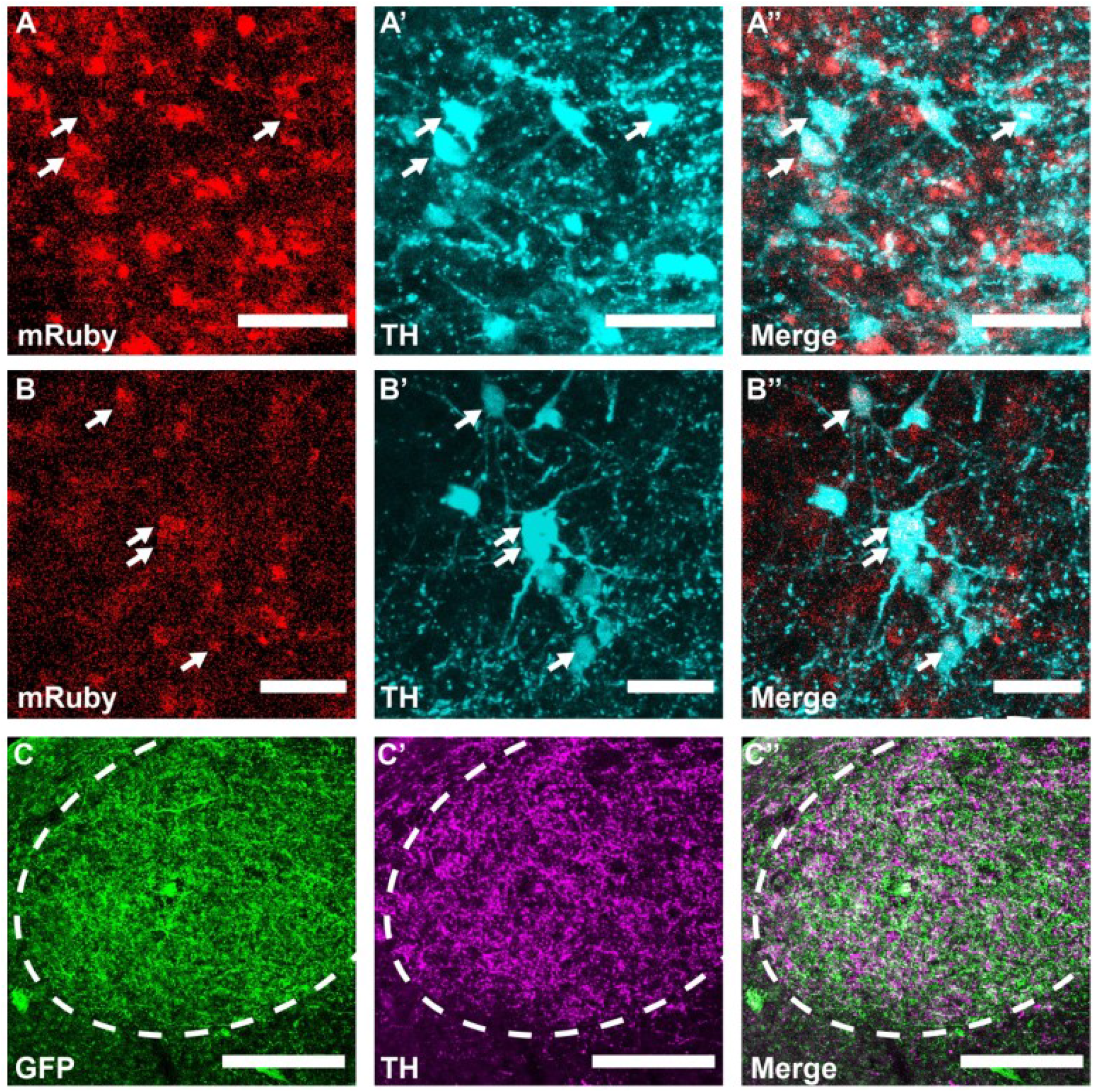
Axon-targeted GCaMP expression in A11 cell bodies and terminals in HVC. (**A**) Representative sagittal section of the injection site in A11 with axon-targeted GCaMP (mRuby, red) and TH antibody labeling (pseudo-colored cyan), scale bars, 50 μm. (**B**) Same as in (A) for a different bird. (**C**) Representative sagittal section of A11 axons in HVC expressing GCaMP (GFP, green) and TH antibody labeling (pseudo-colored magenta), scale bars, 200 μm. Both the GFP and TH labeling is confined to HVC.

**Figure 5 – supplementary figure 2.**
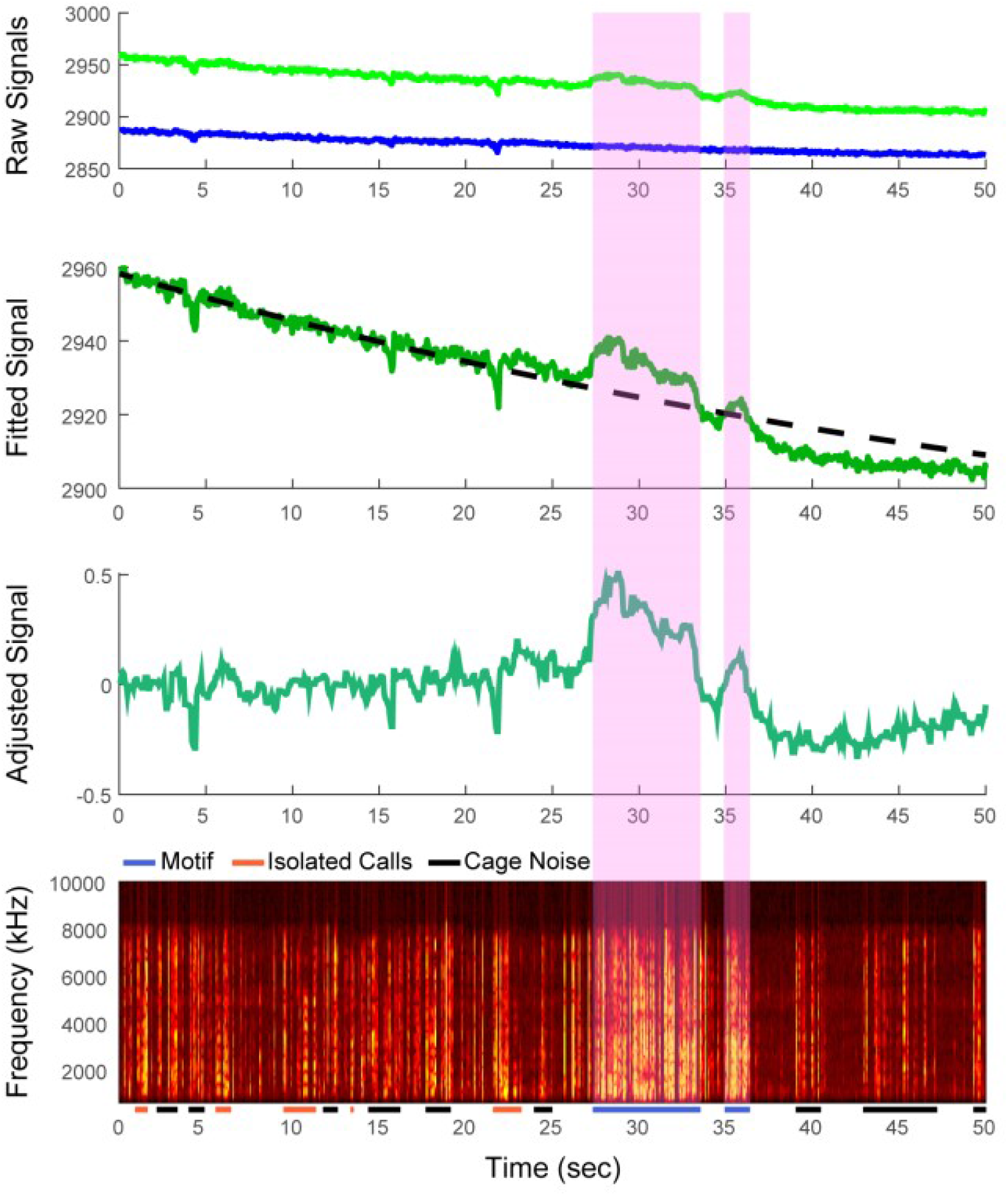
Extraction of A11 axon terminal activity signals in HVC. Top panel: raw signals acquired from exciting HVC with two wavelengths of light: 470nm for imaging of calcium-dependent signals (green) and 415nm for an interleaved isosbestic control (blue). Second panel: regressed control channel (dotted black line) to the trace from the signal channel (green). This regression was used to compute a linear model to generate a predicted signal. Third panel: Filtered calcium-dependent signal after subtracting the predicted control signal from the raw calcium-dependent signal. Bottom panel: spectrogram of the audio recording from the same imaging session. Pink shading denotes instances of female-directed vocalization. Other audio events consist of cage noises and female vocalizations.

**Figure 5 – supplementary figure 3.**
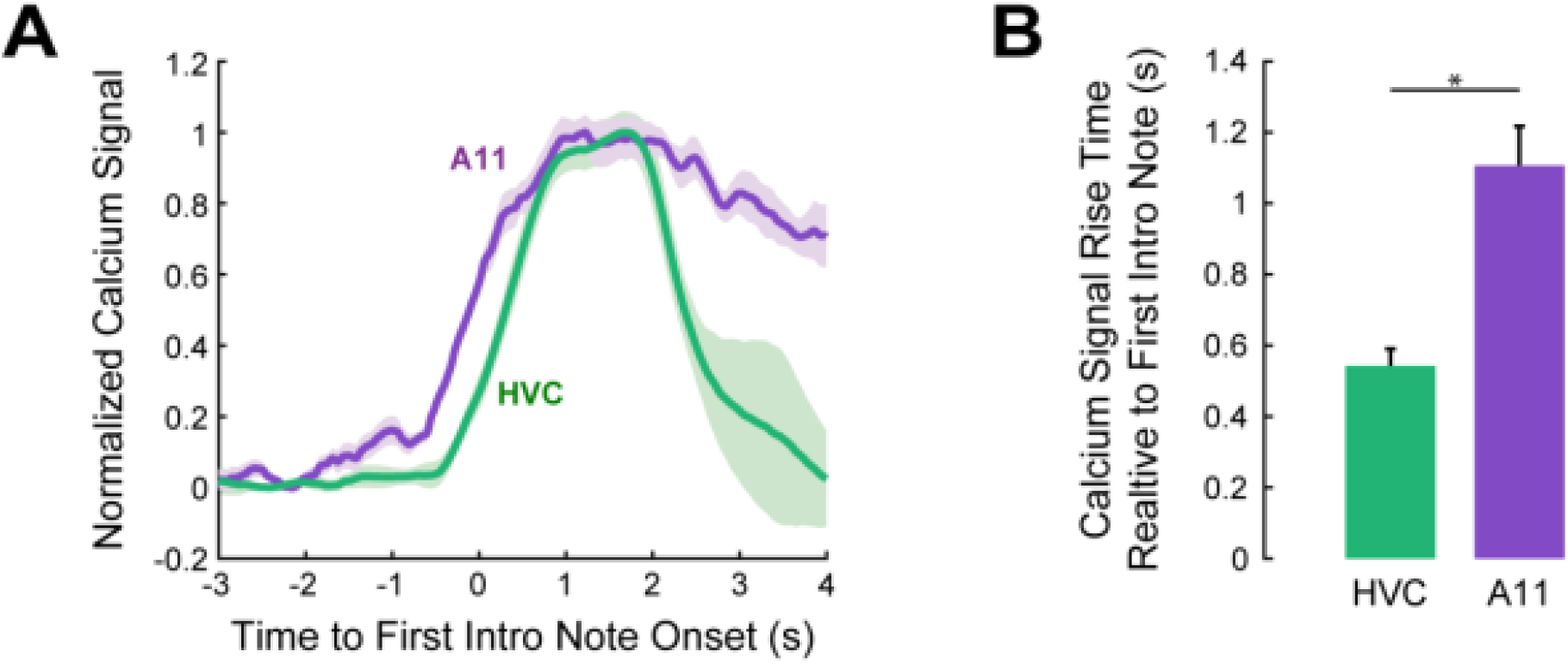
A11 and HVC axons activity aligned to the first introductory notes. (**A**) Calcium traces from one A11 axons bird and one HVC axons bird, aligned to the first introductory note. A11 axons calcium signal start rising before the production of the first introductory note and precedes HVC local axons activity. (**B**) Mean calcium rise time relative to first introductory note for the HVC group (N = 4 birds) and A11 group (N = 4 birds). Calcium rise times for the A11 group is significantly earlier than the HVC group (t-test, p<0.005).

**Figure 5 – supplementary figure 4.**
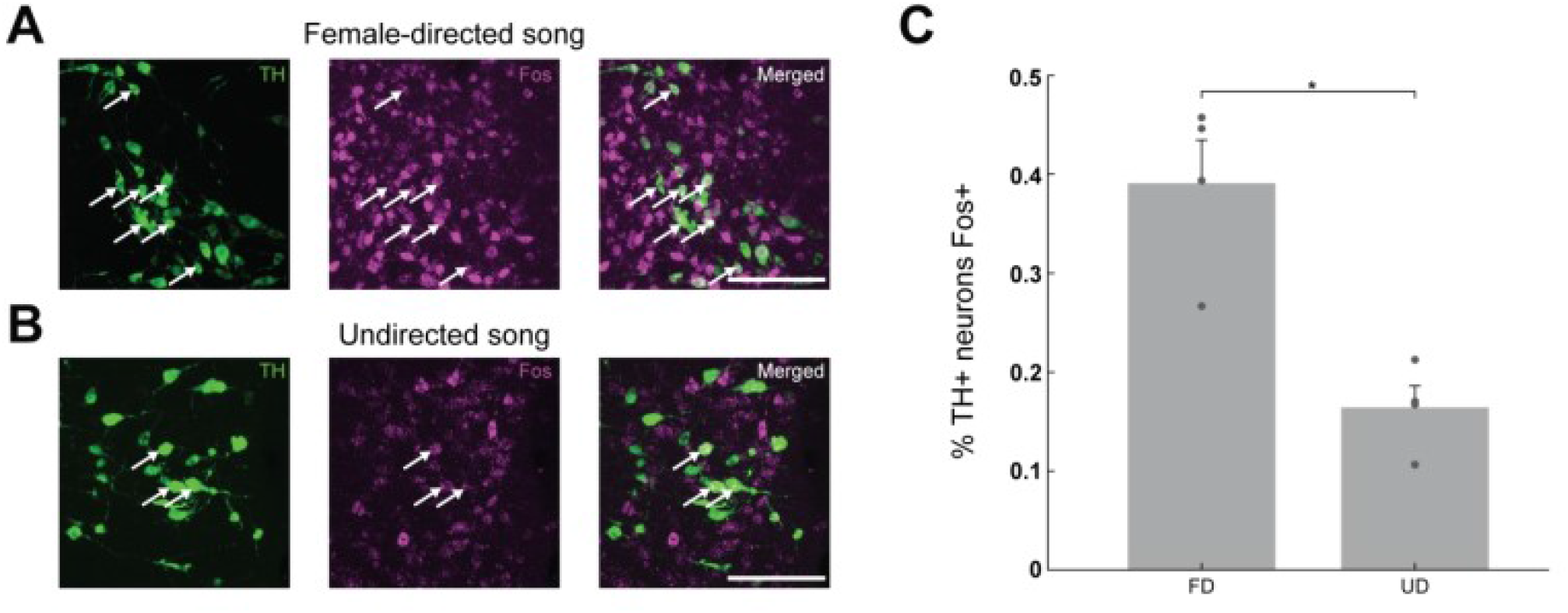
Immunocytochemical labeling of Fos positive A11 neurons. (**A**) TH positive A11 neurons (left panel) Fos positive A11 neurons (middle panel) and double labeled neurons (right panel) of a bird that performed female-directed singing. Scale bar, 100 μm. (**B**) Same as in (A) for a bird that performed undirected songs. (**C**) An elevated Fos response in A11 neurons was shown in males after female-directed song (N=4 birds) relative to males after undirected songs (N = 4 birds) (t-test, p<0.005).

**Movie S1. Social interaction of a naïve male and female zebra finches**

A side view of a male and a female zebra finch with an electronic glass separating between them. When the frame is red, the glass is opaque, and the birds cannot see each other. When the frame turns green., the glass turns clear and the female is visible to the male. The male readily sings to the female as soon as she becomes visible, together with other innate movements. On the bottom: a spectrogram of the audio recorded during the interaction.

**Movie S2. Social interaction of a male after 6-OHDA treatment in HVC with a female**

Social interaction of a male zebra finch that received injections of 6-OHDA into HVC. On the bottom: a spectrogram of the audio recorded during the interaction. Vocalizations consist of repeated introductory notes with no songs.

**Movie S3. Social interaction of a male after 6-OHDA treatment in A11 with a female**

Social interaction of a male zebra finch that received injections of 6-OHDA into HVC. On the bottom: a spectrogram of the audio recorded during the interaction. Vocalizations consist of cage noises.

